# *Clostridioides difficile* PCR ribotype 151 is polyphyletic and includes pathogenic isolates from cryptic clade C-II with mono-toxin pathogenicity loci that can escape routine diagnostics

**DOI:** 10.1101/2022.08.17.504118

**Authors:** Quinten R. Ducarmon, Tjomme van der Bruggen, Céline Harmanus, Ingrid M.J.G. Sanders, Laura G.M. Daenen, Ad C. Fluit, Rolf H.A.M. Vossen, Susan L. Kloet, Ed J. Kuijper, Wiep Klaas Smits

## Abstract

We report a patient case with pseudomembranous colitis associated with a mono-toxin producing *Clostridioides difficile* belonging to the very rarely diagnosed PCR ribotype (RT) 151. The infection was difficult to diagnose, since the isolate and the feces sample tested negative for toxin-encoding genes using a routine commercial test. This prompted us to sequence n = 11 RT151s from various geographical regions to study their genomic characteristics and relatedness. By including whole genome sequence data from other sources, we could further place these isolates into the phylogenetic tree of *C. difficile* and assign them to their respective clades. These analyses revealed that 1) RT151s are polyphyletic with isolates falling into clades 1, and cryptic clades C-I and C-II 2) RT151 contains both non-toxigenic and toxigenic isolates and 3) RT151 C-II isolates contained mono-toxin pathogenicity loci (PaLoc). Additional analysis with PacBio circular consensus sequencing revealed that the isolate from our patient case report contains a novel PaLoc insertion site, lacked *tcdA* and a had significantly divergent *tcdB* sequence that might explain the failure of the diagnostic test. The study is noteworthy as 1) RT151 encompasses both typical and cryptic clades and 2) conclusive evidence for CDI due to clade C-II isolates was hitherto lacking.

## INTRODUCTION

*Clostridioides difficile* is a Gram-positive, spore-forming anaerobic bacterium and the leading cause of antibiotic-associated diarrhea, both in healthcare facilities and the community(1). *C. difficile* can produce several clostridial toxins (toxin A or TcdA, toxin B or TcdB and the binary toxin CDT) and these are responsible for *C. difficile* infection (CDI), symptoms of which can range from mild and self-limiting diarrhea to pseudomembranous colitis, toxic megacolon and ultimately death(1). Phylogenetic analyses have revealed that most *C. difficile* isolates associated with human and animal disease fall within 5 clades(2). In recent years, reports have appeared on (mainly environmental) isolates that are phylogenetically distinct from these 5 clades, but rather fall within at least 3 so-called cryptic clades (C-I, C-II and C-III) that can be considered a separate genomospecies(2-5). Though cryptic clade isolates can contain toxin genes(2, 4, 6), knowledge about the occurrence and clinical significance of these isolates is limited.

Within clades 1-5, the pathogenicity locus harboring the the *tcdA* and/or *tcdB* genes is generally located at a consistent location in the *C. difficile* genome (*cdu1*/*cdd1*) and further includes several other accessory genes (*tcdR, tcdC* and *tcdE*) (6, 7). Importantly, isolates belonging to the cryptic clades C-I and C-III can have divergent pathogenicity locus (PaLoc) sequences and atypical PaLoc insertion sites that can impact diagnostic tests(2, 4, 6, 8).

Here, we report a case of human CDI caused by a RT151 clade C-II isolate for which initial diagnostic tests using Cepheid Xpert *C. difficile* BT assay were negative, even though *C. difficile* could be cultured from patient feces and clinical symptoms corresponded to a CDI case. We show that the cryptic C-II isolate cultured from this patient contains a mono-toxin B PaLoc at a novel location, and that other RT151 isolates fall in phylogenetically divergent clades 1, C-I and C-II.

## MATERIALS AND METHODS

### *Clostridioides difficile* culturing, ribotyping and toxin testing

RT151 isolates analysed as part of this study were derived from various national and international studies in which the Dutch National Expertise Center for *C. difficile* at Leiden University Medical Center (LUMC) participated and had isolates available. The isolates originate from Greece, Malta, The Netherlands and Spain between 2011 and 2021. Detailed information on all RT151 isolates, including relevant accession numbers, sequence types (STs), clade and metadata is provided in Supplemental Table S1.

Isolates were cultured on TSS plates (Tryptic Soy Agar with 5% sheep blood, bioMérieux, The Netherlands) or CLO plates (selective *C. difficile* medium containing cefoxitin, amphotericin B and cycloserine, bioMérieux, The Netherlands). Capillary electrophoresis PCR ribotyping was performed at the Dutch Expertise Centre for *C. difficile*, according to standard procedures(9). Feces samples and *C. difficile* isolates were analyzed by the Xpert *C. difficile* BT assay (Cepheid), that targets *tcdB, cdtA* and a variant *tcdC* gene associated with epidemic strains, according to the manufacturer’s instructions. They were further characterized using an in-house multiplex PCR targeting the 16S rRNA gene, *gluD*, and the genes encoding the large clostrididal toxins and binary toxin(10) or a PCR targeting *tcdB* alone(11). Toxin status was additionally assessed using a toxin neutralization assay using 0.4 μM filter-sterilized culture supernatant from a 48-h culture in Brain-Heart Infusion (BHI) broth on VeroE6 cells, using purified toxins (SML1153-2UG, Lot No SLBT4085, Sigma) and neutralizing anti-TcdA/B antitoxin (T1000, Lot No 1015235, Techlab) (12). Supernatants from 48-h BHI cultures were also analyzed using the VIDAS® *C. difficile* Toxin A & B (Biomérieux), according to the manufacturer’s instructions.

### DNA extraction and whole-genome sequencing

A pure culture from a TSS plate was resuspended in phosphate-buffered saline (Media Products Groningen) to 3 McFarland, and 24 μl of lysozyme (50 g L^-1^) was added to 800 μl of this suspension. The solution was incubated for 20 min at 37°C and 15 μl of proteinase K (20 g L^-1^l; Roche) was added, followed by a 20 min incubation at 37°C. DNA was extracted on the QiaSymphony platform (Qiagen, The Netherlands), using the Qiasymphony Virus/Pathogen Midi Kit according to the manufacturer’s instructions. Isolated total DNA was submitted for sequencing to Genomescan (Leiden, The Netherlands) on the Illumina Novaseq6000 platform with read length 150 bp in paired-end mode.

To generate complete genomes, three isolates (LUMCMM21 0001/0003, LUMCMM19 2333 and LUMCMM16 0013) were submitted for long read sequencing on the Pacific Biosciences (PacBio) Sequel platform at the Leiden Genome Technology Center. To prepare high molecular weight total DNA, isolates were cultured overnight in 10 mL BHI at 37°C. Cells from 5 mL of culture were pelleted and processed using the Qiagen Genomic-tip 100/G, according to the manufacturer’s instructions. PacBio sequencing libraries were generated as follows. Genomic DNA was sheared with the Megaruptor 3 system (Diagenode) using 35-40 cycles. SMRTbell® libraries were generated according to the following manufacturer’s procedure & checklist: Preparing whole genome and metagenome libraries using SMRTbell® prep kit 3.0, PN 102-166-600 APR2022. To retain DNA corresponding to plasmids <10 kb, no size-selection was applied. The libraries were sequenced on a PacBio Sequel II platform with a 30 h movie time using Binding Kit v2.2, Sequencing Primer v5 and Sequencing Kit v2.0.

### *C. difficile* genome retrieval

Four additional RT151 genome sequences were obtained from the laboratory collection of the UK Anaerobic Reference Unit (Supplemental Table S1); these were processed and sequenced previously as part of their routine surveillance and typing activities but raw Illumina reads were processed in parallel with the other RT151 genomes reported here for the purpose of this study, as described below.

To assess the phylogenetic relationship of the eleven RT151 in the context of the previously established clades of *C. difficile*, we downloaded additional isolate sequence data: we included two isolates of each of the three cryptic clades (C-I, C-II and C-III)(2), 4 isolates of clade 1 (as this is the largest group) and two isolates of clade 2-5 from the Leiden-Leeds collection(13, 14) (Supplemental Table S1).

### Data processing, assembly and quality control

Raw Illumina sequence data was cleaned and trimmed using fastp (v0.23.2)(15) and sequence quality was inspected using Fastqc (v0.11.9) and Multiqc (v1.8)(16, 17). Cleaned reads were assembled by using SKESA (v2.4.0) and SPAdes (v3.15.3) with “–untrusted-contigs” and “–isolate” parameters(18, 19). Only contigs with length 500 >= bp were kept. Assembly quality and general characteristics of the assemblies were assessed using QUAST (v5.0.2)(20). The scaffolds obtained from SPAdes were used for downstream analyses. Default parameters were used unless stated otherwise.

Raw PacBio sequence data from LUMCMM16 0013 (RT151, clade C-I), LUMCMM21 0001 (RT151, clade C-II) and LUMCMM19 2333 (RT151, clade C-II) was assembled using Flye (v2.9)(21)” and the start position was fixed using Circlator and the “fixstart” parameter(22). Assembly quality was subsequently inspected using available output metrics from Flye, QUAST and completeness using BUSCO (v5.3.2, dataset clostridia_odb10 creation date 2020-03-06)(23). Analysis of m4C and m6A methylation was performed using PacBio SMRT® Link Software (v11.0.0.146107).

### *C. difficile* genome annotations and sequence typing

Prokka (v1.14.6) was used for rapid genome annotation and to obtain protein sequences, with the “– kingdom Bacteria”, “--genus Clostridioides” and “–species difficile” parameters(24). For pan-genome and core-genome analyses, Roary (v3.13.0) was ran on the PROKKA-annotated scaffolds with the “-e mafft” parameters(25, 26).

The accessory genome was subsequently visualized in R (v4.2.0) using the ComplexHeatmap package (v2.12.0)(27). To construct a phylogenetic tree based on the core genes as identified by Roary, we used IQTree (v2.2.0) with the “-B 2500” and “-m MFP” parameters for 2500 bootstrap replicates and automated model selection(28, 29). The best model was chosen based on the minimized BIC score and was identified to be UNREST+F0+I+G4. The obtained phylogenetic tree was subsequently visualized using iTOL v6 and the tree was rooted at midpoint(30).

Multi-locus sequence typing (MLST) was performed using mlst (v2.19.0) with the PubMLST *C. difficile* database updated to October 21^st^ 2021(31). To assign clades to each of our sequence types, we used the information contained in Knight et al, which includes a clade assignment for sequence types(2). Novel loci (sodA-116, recA-82 and glyA-140) were identified and their respective sequences obtained by mlst using the “-q –novel parameters” and submitted to the pubmlst database. Based on this, three isolates were assigned new sequence types (ST946, ST947 and ST948) through PubMLST (https://pubmlst.org/). Lastly, pairwise average nucleotide identities (ANIs) between the genomes were calculated using the fastANI tool (v1.33)(32).

Available metadata for all isolates, including newly assigned sequence types, are listed in Supplemental Table 1.

## Data availability

Raw sequence data is available at the European Nucleotide Archive under BioProject PRJEB52887. Associated metadata can be found in Supplemental Table S1. All bioinformatic tools used for the analyses are freely available through the references provided. Complete chromosomes for PacBio-sequenced isolates can be found as accession numbers GCA_945861085 (LUMCMM21 0003), GCA_945909465 (LUMCMM19 2333) and GCA_945909635 (LUMCMM16 0013). For PaLoc alignment and insertion analyses, the corresponding regions were extracted from the complete genome sequence of the reference strain *C. difficile* 630 (AM180355)(33). The pHSJD-312 sequence was obtained from accession MG973074(4). The patient provided written informed consent for the anonymized materials and results to be used for publication.

## RESULTS AND DISCUSSION

### Case description and identification of a RT151 isolate from a CDI patient

In 2021, a 46-year-old female patient was diagnosed with acute myeloid leukemia and treated with two courses of induction chemotherapy followed by allogeneic hematopoietic stem cell transplantation (allo-HSCT) at the University Medical Center Utrecht (UMCU), a tertiary care center in the Netherlands. Due to primary graft failure, the patient received a second allo-HSCT two months later. Between the first allo-HSCT and engraftment after the second allo-HSCT, the patient was neutropenic for a prolonged time. This period was complicated by candidemia with *Candida kefyr*, treated with anidulafungin followed by fluconazole, two bacteraemia episodes with a multidrug resistant *Klebsiella pneumoniae* treated with imipenem/cilistatin and probable pulmonary aspergillosis for which the patient received antifungal treatment. The post-transplantation period was complicated by early reactivation of cytomegalovirus treated with foscarnet and later letermovir prophylaxis, and mild graft-versus-host disease of the skin for which the patient received topical steroids.

In the week prior to the second allo-HSCT, the patient developed diarrhea (day 0) and feces was positive for norovirus and sapovirus as tested by PCR. The commonly used molecular test for *Clostridioides difficile* (Cepheid Xpert *C. difficile* BT) was negative on a stool sample at day 0. Despite the use of loperamide, diarrhea persisted in the following weeks although subsequent PCRs for noro-and sapovirus became negative. PCRs for rotavirus, enterovirus, *Salmonella, Shigella, Yersinia, Campylobacter, Plesiomonas, Giardia, Cryptosporidium, Dientamoeba fragilis, Blastocystis hominis, Entamoeba histolytica* were also negative. The Xpert *C. difficile* BT assay was repeatedly negative on day 43, day 45 and day 49. Colon biopsies taken on day 53 showed no signs of graft-versus-host disease or pseudomembranous colitis. Around day 50 to 55, diarrhea seemed to diminish, but around day 65 it started to intensify with up to ten watery stools per day in the ensuing weeks.

Repeated stool tests for viral, bacterial and parasitic infection were negative, including microscopy for helminth eggs, cysts and *Cyclospora* and the Xpert *C. difficile* BT (on day 66, day 80, day 84 and day 86). However, macroscopic findings during colonoscopy performed on day 82 were suggestive of pseudomembranous colitis, confirmed by microscopy of colon biopsies while there were no signs of graft-versus-host disease. These findings prompted treatment with oral vancomycin starting on day 84 using a taper-and-pulse regimen. After 4-5 days, the patient started to improve considerably with stools becoming less frequent, lower stool volumes and a more solid consistency. Ethanol-treated fecal material obtained on day 84 was culture-positive for *C. difficile* on cycloserine cefetoxin fructose agar after 48h anaerobic incubation, as confirmed by matrix-assisted laser desorption/ionization time-of-flight mass spectrometry (MALDI-TOF score of 2.09, database BDAL-8468, Bruker Daltonics, Germany). On day 159 (35 days after completion of anti-CDI treatment), the patient developed a mild episode of diarrhea with abdominal cramps at home, with negative Xpert *C. difficile* BT test on stool but again a positive culture of *C. difficile* from feces (MALDI-TOF score of 2.05) that was later confirmed to belong to RT151. The patient spontaneously recovered without antibiotic treatment within the two weeks that followed. Both isolates tested negative in the Xpert *C. difficile* BT assay, but positive when using an in–house developed PCR for *C. difficile* toxin B(11). Both isolates were then sent to the Dutch Expertise Centre for *C. difficile* at the LUMC, with the first isolate receiving the identification number LUMCMM21 0001 and the second isolate LUMCMM21 0003. Both isolates were characterized by capillary PCR ribotyping at the LUMC and were shown to be RT151.

### RT151 isolates have different toxin profiles

To obtain a better insight into the prevalence and characteristics of RT151, we retrieved all isolates from the collection of the Dutch Expertise Center for *C. difficile*, that encompasses > 22000 isolates collected since 2005. Seven isolates had been typed as RT151 using either agarose-gel based ribotyping or capillary electrophoresis-based PCR ribotyping, demonstrating that this is a very rarely diagnosed PCR ribotype (<0.035%).

Within this collection of RT151 isolates, five were non-toxigenic *C. difficile* (NTCD) and two were toxigenic on the basis of a multiplex PCR. The isolate obtained from the case description from Utrecht (LUMCMM21 0001) was positive for the gene encoding toxin B, but not the gene encoding toxin A (A^-^B^+^), while an isolate obtained from a patient in Malta in 2019 (LUMCMM19 2333) had a mono-toxin A (A^+^B^-^) PaLoc and neither isolate contained the binary toxin locus according to PCR results.

These results are in line with findings of the Leeds Teaching Hospitals NHS Trust and University of Leeds that report a prevalence for RT151 of <0.03% with varying toxin status in a cell cytotoxicity and neutralization assay (CCNA; personal communication K.A. Davies and J. Freeman). In addition, the global *C. difficile* analysis carried out by Knight and colleagues also found a prevalence of 0.033% for ST205, based on a collection of 12,098 genomes(2).

We also assessed toxigenicity using a diagnostic enzyme-linked fluorescent assay (VIDAS®, bioMérieux) and a VeroE6-based CCNA. The result showed that only the *tcdB*-positive LUMCMM21 0001/0003 tested positive for toxin production (Supplemental Table S1). No toxin production could be demonstrated for the *tcdA*-containing isolate (LUMCMM19 2333) under the conditions tested.

### Core genome phylogeny places some RT151 isolates in cryptic clade C-I and C-II

To determine the relatedness between our RT151 isolates and to understand their placement in the phylogenetic tree of *C. difficile*, we performed Illumina short-read whole genome sequencing (WGS). In addition to the seven RT151 isolates obtained from our in-house collection, the UK Anaerobic Reference Unit provided WGS data from four additional RT151 isolates. From a detailed analysis it became clear that the two isolates isolated from the patient described above (LUMCMM21 0001 and LUMCMM21 0003) are identical (data not shown).

By placing the eleven RT151 isolates into a phylogenetic tree with representative isolates from cryptic clades I-III clades and all ‘classical’ clades to ensure genetic diversity in our tree, we assigned the RT151 isolates to their respective clades based on 1158 core genes in our analysis (Figure 1). This analysis revealed that the majority (n = 8) of our RT151 isolates belonged to Clade 1 and MLST typing revealed these to be ST205 (or highly related ST type(s); ST205-like) (Supplemental Table 1 and Figure 1). All of the RT151 isolates obtained from the UK Anaerobic Reference Unit fell into ST205 (or ST205-like) groups and were therefore not further analyzed. Interestingly, three of the isolates were classified into cryptic clades, with two independent isolates belonging to clade C-II (LUMCMM21 0001/0003 and LUMCMM19 2333) and one isolate to clade C-I (LUMCMM16 0013). To complement the phylogenetic analysis based on core genes, we next computed ANIs of our cryptic C-II isolates versus all other genomes (Fig 1B and C). This analysis shows that our isolates have ANIs of > 98.7% against all other C-II isolates, but that ANI values to other clades fall well below the 96% that has been named as a cut-off for the same species(34).

**Figure 1.**
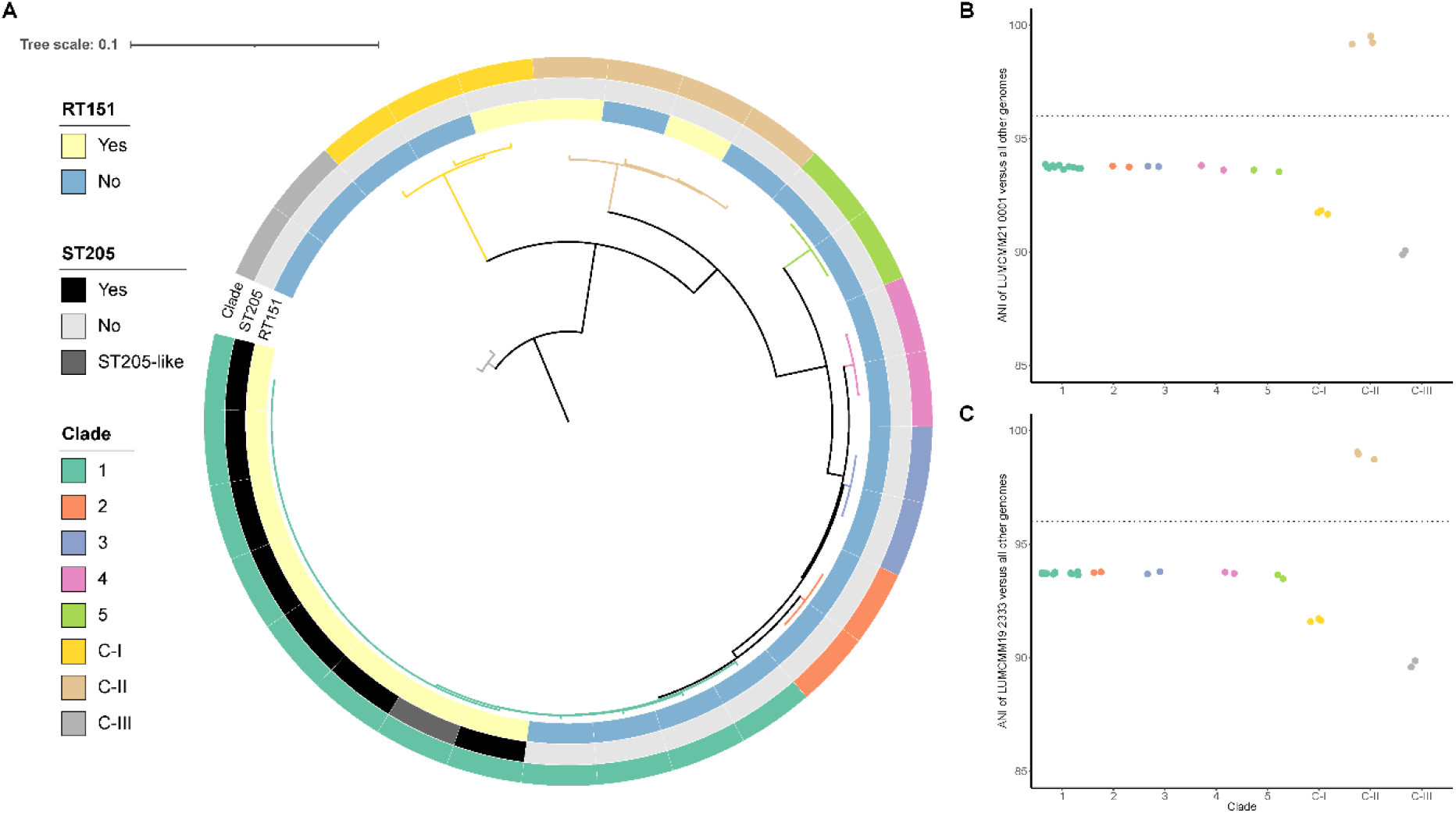
RT151 isolates are part of clade 1, cryptic clade C-I and C-II and belong to various sequence types. **A**. The tree was rooted at midpoint in iTOL version 6. Detailed information, including the depicted classifications, are available in Supplemental Table S1. **B**. ANI calculations of LUMCMM21 0001/0003 versus all other included isolate genomes. **C**. ANI calculations of LUMCMM19 2333 versus all other included isolate genomes.

These data demonstrate that *C. difficile* ribotype 151 is polyphyletic with isolates from the same ribotype falling into two different cryptic clades as well as a ‘regular’ clade, which has not been reported before. Though there is good concordance between PCR ribotype and MLST-based approaches(13, 35, 36), these findings underscore that PCR ribotyping is insufficiently discriminatory for phylogenetic placement.

### A pan-genome analysis suggests that accessory genes are conserved within ST205

It is conceivable that despite divergent core genomes, RT151 share a common accessory genome. Therefore, we next investigated the accessory genes. We identified a total of 14,297 accessory genes and these were clustered using the complete linkage method with Euclidian distances in the ComplexHeatmap package(27) to identify potential similarities in the accessory genome of RT151 or ST205 (Figure 2). This analysis largely confirms the core genome phylogeny (Figure 1), as a clear clustering based on accessory genes is seen within RT151 isolates belonging to ST205, while the cryptic clade isolates from RT151 have divergent accessory genomes. In addition, isolates from the same clade also generally cluster together, with the exception of clade 2 isolates falling within the larger clade 1 cluster.

**Figure 2.**
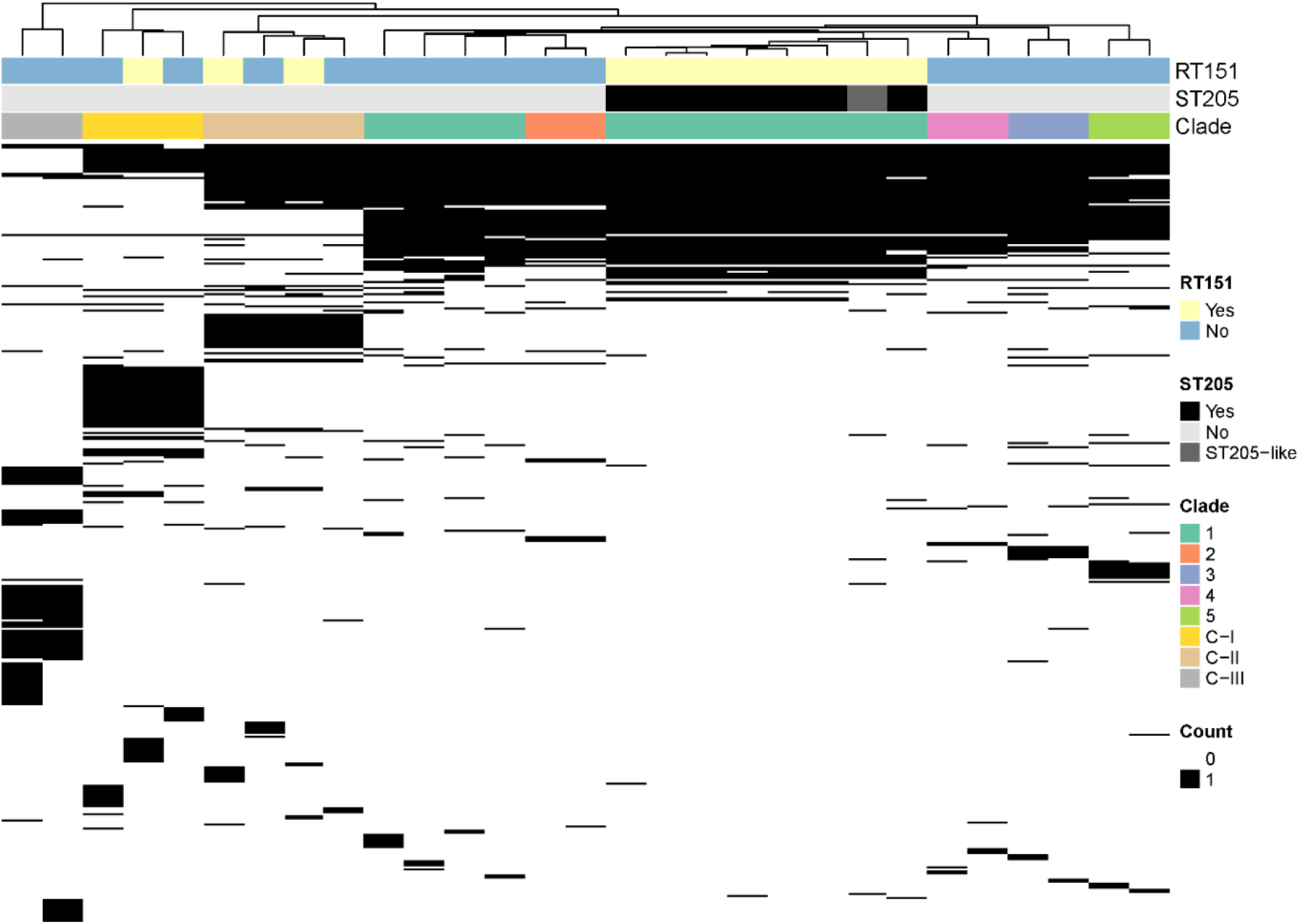
No clustering of RT151 isolates based on the accessory genome (n = 14,297 genes). Black denotes the presence of a gene, white denotes the absence of a gene. Clustering of accessory genes was performed using the complete linkage method with Euclidian distances in the ComplexHeatmap package(27). Detailed information, including the depicted classifications, are available in Supplemental Table S1. Coloring is the same as in Figure 1.

### Cryptic clade C-II RT151 isolates can have a mono-toxin pathogenicity locus at a novel insertion site

On the basis of our initial Illumina-sequenced genomes we were unable to recover PaLoc sequences on a single contig for all isolates and, importantly, genomic neighborhoods around the PaLoc could not be fully resolved. In particular, for the *tcdB*-positive C-II isolate from the Netherlands (LUMCMM21 0003), *tcdB* was contained on a 18-kb contig lacking flanking chromosomal regions. Additionally, we noted limited homology to the pHSJD-312 plasmid (Supplemental Figure S2A)(8), which carries a similar PaLoc. However, in contrast to the PaLoc on this plasmid, the LUMCMM21 0003 PaLoc lacked the binary toxin genes, *cdtA* and *cdtB*.

To identify the PaLoc insertion site and to establish whether it is carried on plasmid, we generated complete genomes for the three cryptic clade RT151 isolate using PacBio circular consensus sequencing (Table 1). The non-toxigenic C-I isolate (LUMCMM16 0013) was found to contain extrachromosomal elements (ECEs) of 132 kb and 60 kb, that are classified in a PHASTER prediction as possible and putative phage, respectively. The *tcdB* positive C-II isolate (LUMCMM21 0003) showed only a single contig, demonstrating that the toxin gene is not carried on a plasmid. The *tcdA*-positive C-II isolate (LUMCMM19 2333) harbors a 13-kb ECE plasmid but this plasmid does not contain toxin genes. Thus, in contrast to previous reports from C-I strains(4, 8), the cryptic RT151 isolates appear to carry their toxins on the chromosome. To further classify our toxin genes, we used DiffBase(37) and this revealed LUMCMM21 0003 to have tcdB subtype B12, while LUMCMM19 2333 had toxin subtype A7.

**Table 1.**
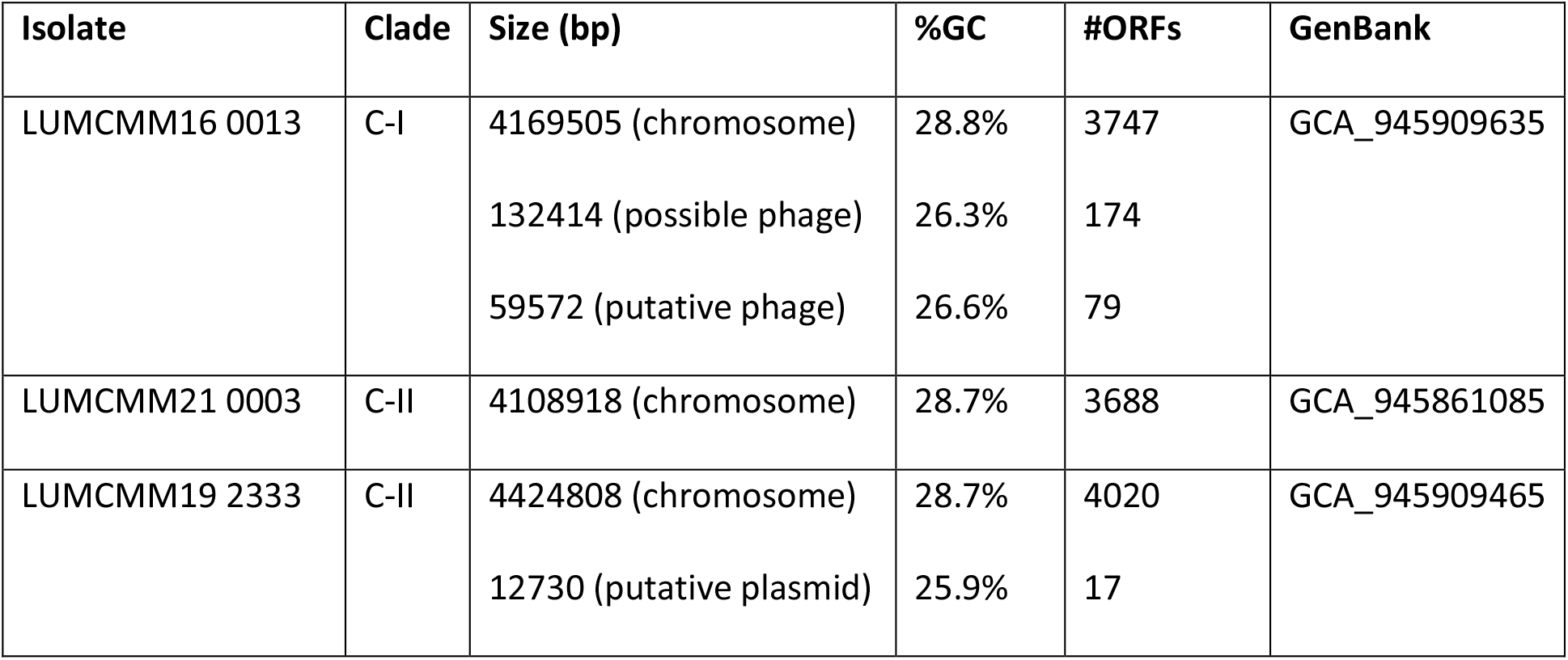
Genome characteristics of the cryptic clade RT151 isolates.

PacBio sequencing also allows us to identify motifs associated with m4C and m6A methylation events (Supplemental Table S2). Methylation has been reported to affect virulence characteristics of *C. difficile*(38). Notably, all strains contain the CamA-dependent modification of CAAAA**A** sequences (modified residue in bold)(38, 39), indicating that the action of this methyltransferase is conserved in the RT151 C-I and C-II strains. We also observed m6A methylation on a subset of CAAA**A**AASNV motifs in the C-II strains, but not the C-I strain, suggesting it may be specific for clade C-II. Finally, we observed m4C modification on a subset of **<u>C</u>**TATTATCW motifs in the C-II strains, that may be conserved in C-I strains (CWATTATCW). This motif has not previously been identified by others(38).

Next, we investigated the genomic location of the PaLoc’s of the RT151 isolates by aligning the observed insertion sites to those known ones from literature (Figure 1A)(4, 6), using the genome sequence of the RT012 reference strain 630 and the sequence of plasmid pHSJD-312as a reference (Figure 3)(4, 33). This analysis reveals that the PaLoc in the Dutch C-II isolate is inserted at a location that has not previously been described, between genes homologous to *cd0628* and *cd0629* of strain 630 (Figure 3C).

**Figure 3.**
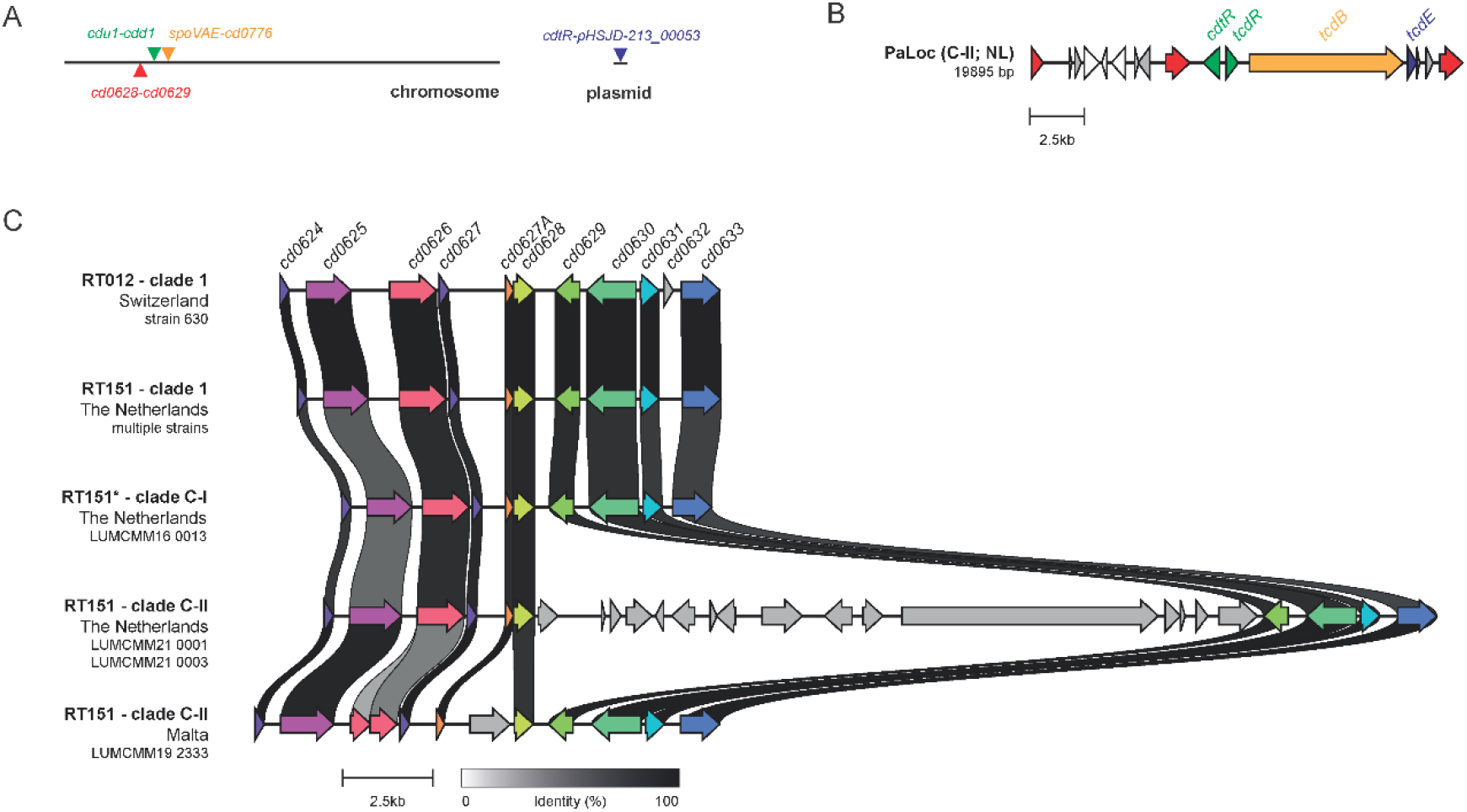
Structure of the mono-toxin pathogenicity locus and PaLoc insertion sites of two cryptic clade-II RT151 strains. **A**. Overview of known PaLoc insertion sites (above the lines)(4, 6) and the novel PaLoc insertion site (below the line) identified in the present study. Three sites are located on the chromosome (nomenclature from RT012 reference strain 630)(33), and one is located on a plasmid (nomenclature from clade C-I strain strain HSJD-312)(4, 8). **B**. Structure of the PaLoc in the A-B+ RT151 clade C-II strain (LUMCMM21 0001/0003) showing all open reading frames predicted by Prokka(24). Genes involved in transposition or recombination are indicated in red, hypothetical genes with homology to the PaLoc on pHSJD-312 are indicated in gray, genes encoding toxin regulators are indicated in green, the toxin gene is in orange and the holin gene in purple. **C**. Alignment of the chromosomal region incorporating the *cd0628*-*cd0629* insertion site for RT151 isolates in comparison with the RT012 reference strain 630(33). RT151 clade 1 is LUMCMM11 0004, RT151 clade C-I is LUMCMM16 0013, and RT151 clade C-II are LUMCMM21 0001/0003 (The Netherlands) and LUMCMM19 2333 (Malta). Figure generated by clinker(43)and centered on *cd0628*. Colored arrows indicate similar genes; links are drawn between similar genes on neighboring clusters and are shaded based on sequence identity (0% white, 100% black, identity threshold for visualization 0.35).

The PaLoc of LUMCMM21 0003, as well as the *cdu1*-*cdd1* insertion site that normally harbors the PaLoc, is reminiscent of what has been described for a toxin A-negative toxin B-positive toxinotype XXXII strain, 173070(5). Similar to our isolate, the *cdu1*-*cdd1* locus contains a multi-gene insertion interrupted by an IS256 insertion element and the PaLoc has a *tcdR*-*tcdB*-*tcdE* architecture, lacks the binary toxin genes but has a divergently transcribed transcriptional regulator gene upstream from *tcdR* (presumably this is *cdtR*). By downloading raw reads (SRR1514909) and performing ANI analyses on the assembled genome, we confirmed that 173070 was also a cryptic C-II isolate (data not shown). This is in line with the SLO148 ribotype reported in the manuscript(5) to be highly similar to that of RT151 (data not shown) and the reported ST (ST200) is classified as C-II in other studies(2, 40).

The PaLoc in the Maltese C-II isolate (LUMCMM19 2333) is inserted between *spoVAE* and *cd0776*, as previously described for isolate RA09-70(6) (Figure 1A, Supplemental Figure S2). RA09-70 was classified as Clade 5 but the phylogenetic analysis by the authors of the manuscript included only strains from clade 1-5 and C-I. We therefore downloaded the respective genome assembly (GCF_001299495.1) and confirmed through ANI analyses that RA09-70 is, in fact, a clade C-II isolate that was misclassified as clade 5 for lack of reference (data not shown).

### The role of C-II isolates in CDI

To the best of our knowledge, the present study is the first to conclusively link a clade C-II isolate to human disease, although the above sentences indicate that clade C-II isolates could be more common than hitherto assumed. Nevertheless, diagnosis of CDI due to LUMCMM21 0001/0003 was hampered by the fact that fecal samples, as well as *C. difficile* isolates, tested negative in the Xpert *C. difficile* BT assay, used by many laboratories worldwide. Toxin gene sequences of cryptic clade isolates can differ significantly from those of regular clade 1-5 isolates which can lead to false negative results(4, 5). Due to the proprietary nature of the Xpert *C. difficile* BT assay, we could not confirm that this failure is due to mismatches between the primers used in the assay and the C-II *tcdB* sequence but we have informed the company and provided the WGS data to investigate this further. We do note that positive identifications were made using the ECDC-endorsed multiplex PCR (relating to little or no mismatches in the primer sequence compared to the genome). A recent report investigating *C. difficile* isolates from environmental locations showed several atypical PaLoc locations in cryptic clade isolates and *in silico* PCR further suggested that routine diagnostic assays would return highly variable results if such isolates would be encountered in the clinic(3), which is in line with what we found in the current study. This underscores the potential of these isolates to be OneHealth pathogens. Toxin-based assays (CCNA and VIDAS®) demonstrated toxin production for the mono-toxin B isolate (LUMCMM21 0001/0003), but not the mono-toxin A isolate (LUMCMM19 2333). Though the latter was isolated from a hospitalized patient, lack of metadata did not allow us to establish whether the patient had symptoms consistent with CDI and it remains to be established if this isolate is pathogenic. Recently, it has been reported that toxin A-positive toxin B-negative C-III strains cannot be identified on ChromID agar due to their esculin-negative nature(41). We found, however, that all RT151 strains – including those belonging to clades C-I and C-II, were esculin-positive (Supplemental Table S1).

Altogether, reports of cryptic clade isolates that cause human CDI are still very rare. With the increasing rate of community-onset CDI(42) and the discovery that cryptic clade isolates with atypical PaLoc configurations are widespread in the environment (3), it is very likely that other CDI cases caused by isolates from this cryptic clade occur but are missed by standard diagnostics if only PCR is used.

## Acknowledgements

The authors thank K. Galea (L-Università ta’ Malta); J. Freeman and K. Davies (University of Leeds) for sharing unpublished data; Michael Perry and Trefor Morris (Anaerobic Reference Unit, Cardiff) for providing RT151 whole genome sequence information; I. Sidorov for implementation of clinker; and members of the Experimental Bacteriology Group at the LUMC for helpful discussions.

## Author contributions

QRD: software, validation, formal analysis, data curation, writing - original draft, visualisation; JTvdB: writing – original draft; CH: investigation; IMJGS: investigation; LD: resources; ACF: investigation; RHAMV: investigation, formal analysis; SLK: resources, supervision; EJK: conceptualization, writing - original draft; WKS: conceptualization, formal analysis, writing - original draft, visualization, supervision. All authors reviewed and edited the manuscript, and approved the final version.

## SUPPLEMENTAL INFORMATION

**Supplemental Table S1.**
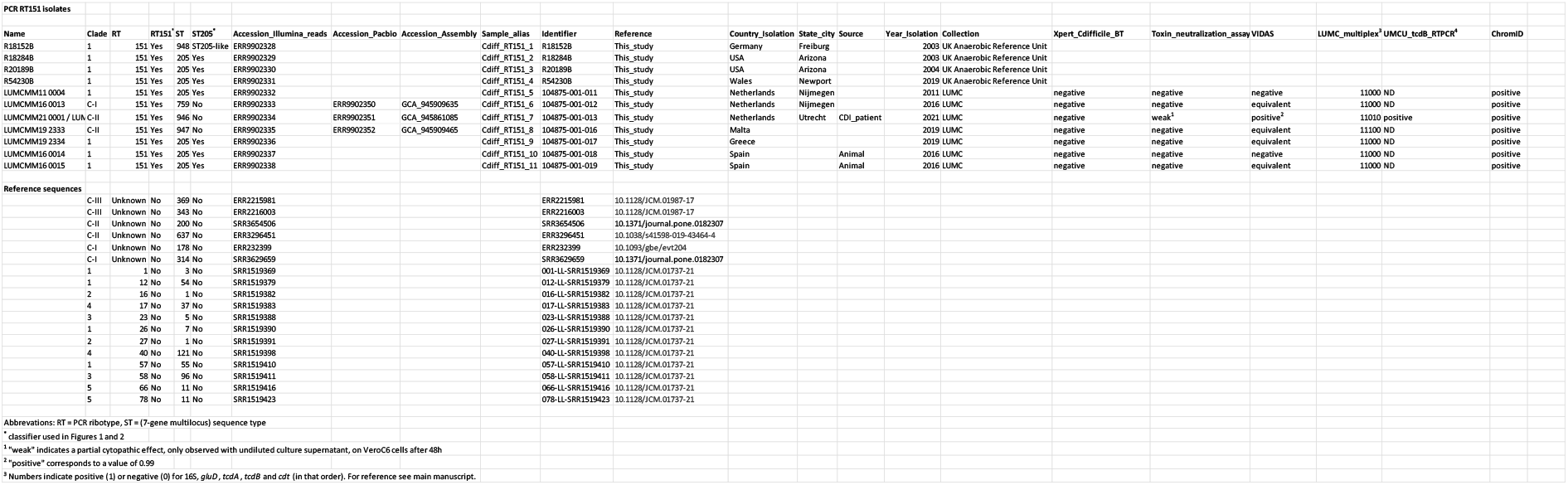
Characteristics of RT151 isolates and reference sequences analyzed in this study.

**Supplemental Table S2.**
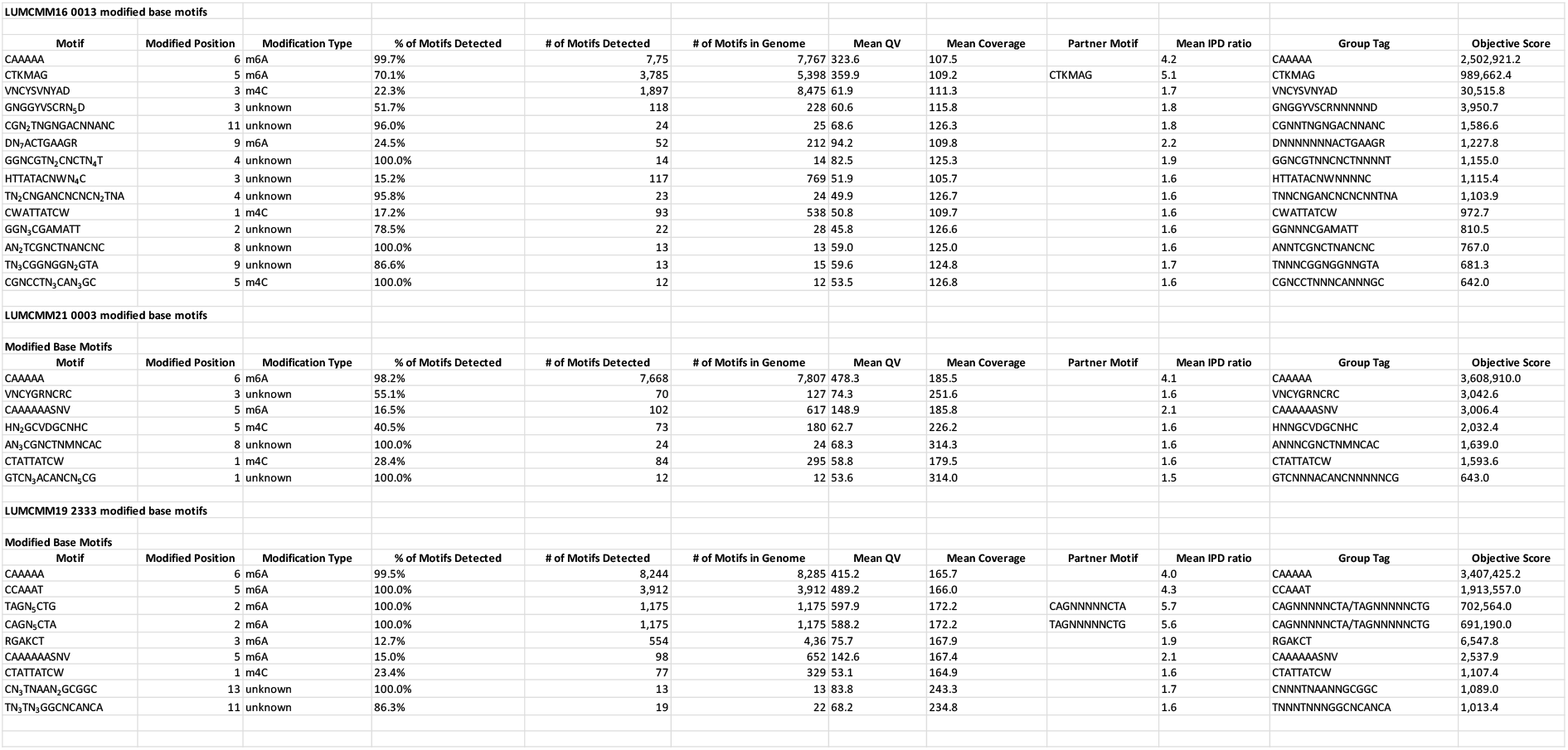
Overview of modified base motifs of the cryptic clade RT151 isolates.

**Supplemental Table S3.**
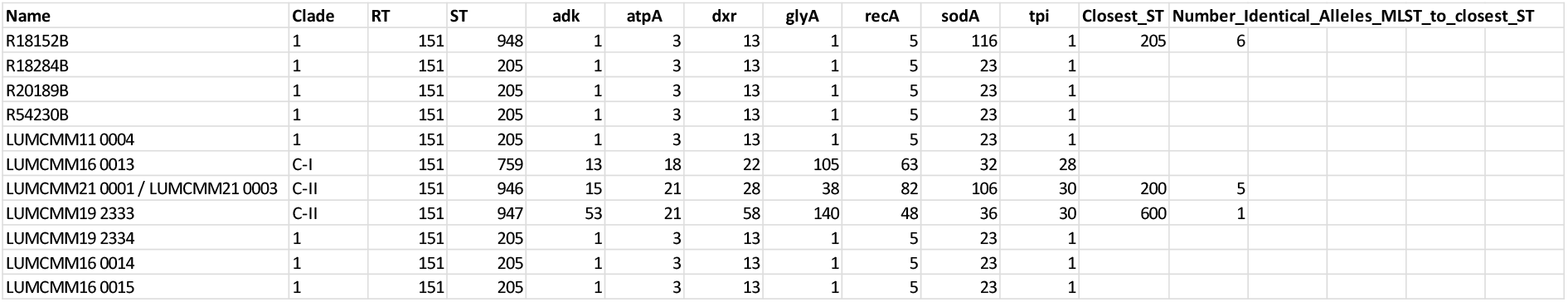
Overview of MLST profiles of all RT151 isolates.

**Supplemental Figure S1.**
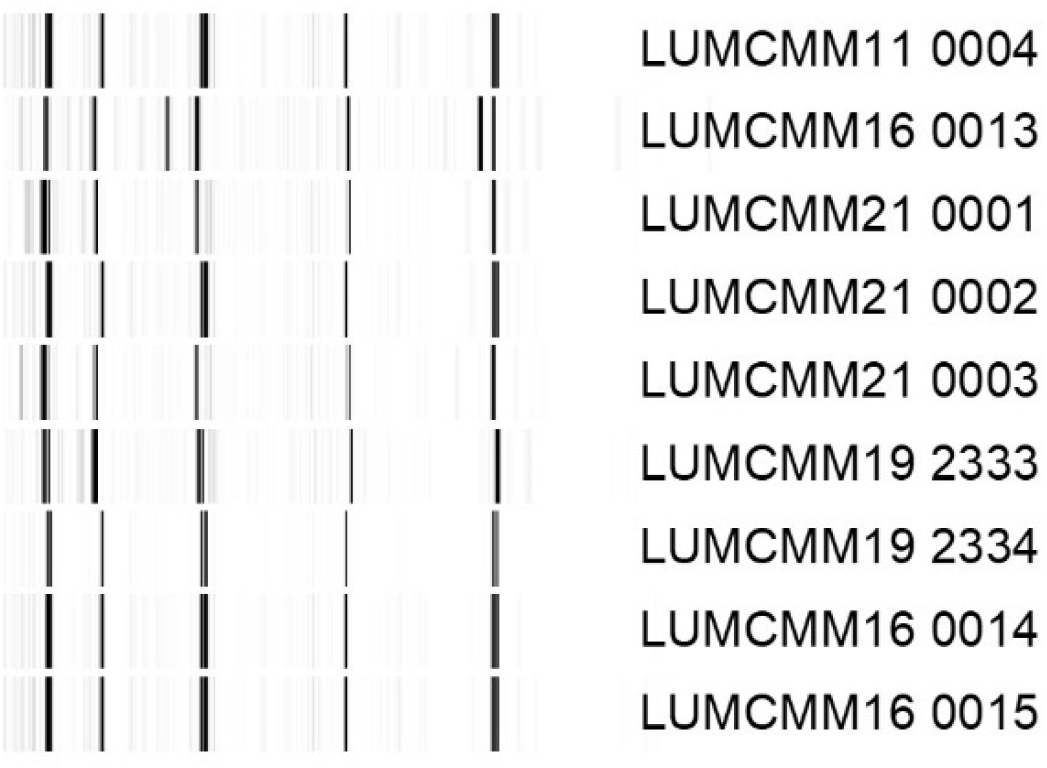
PCR RT banding profiles of all RT151 isolates.

**Supplemental Figure S2.**
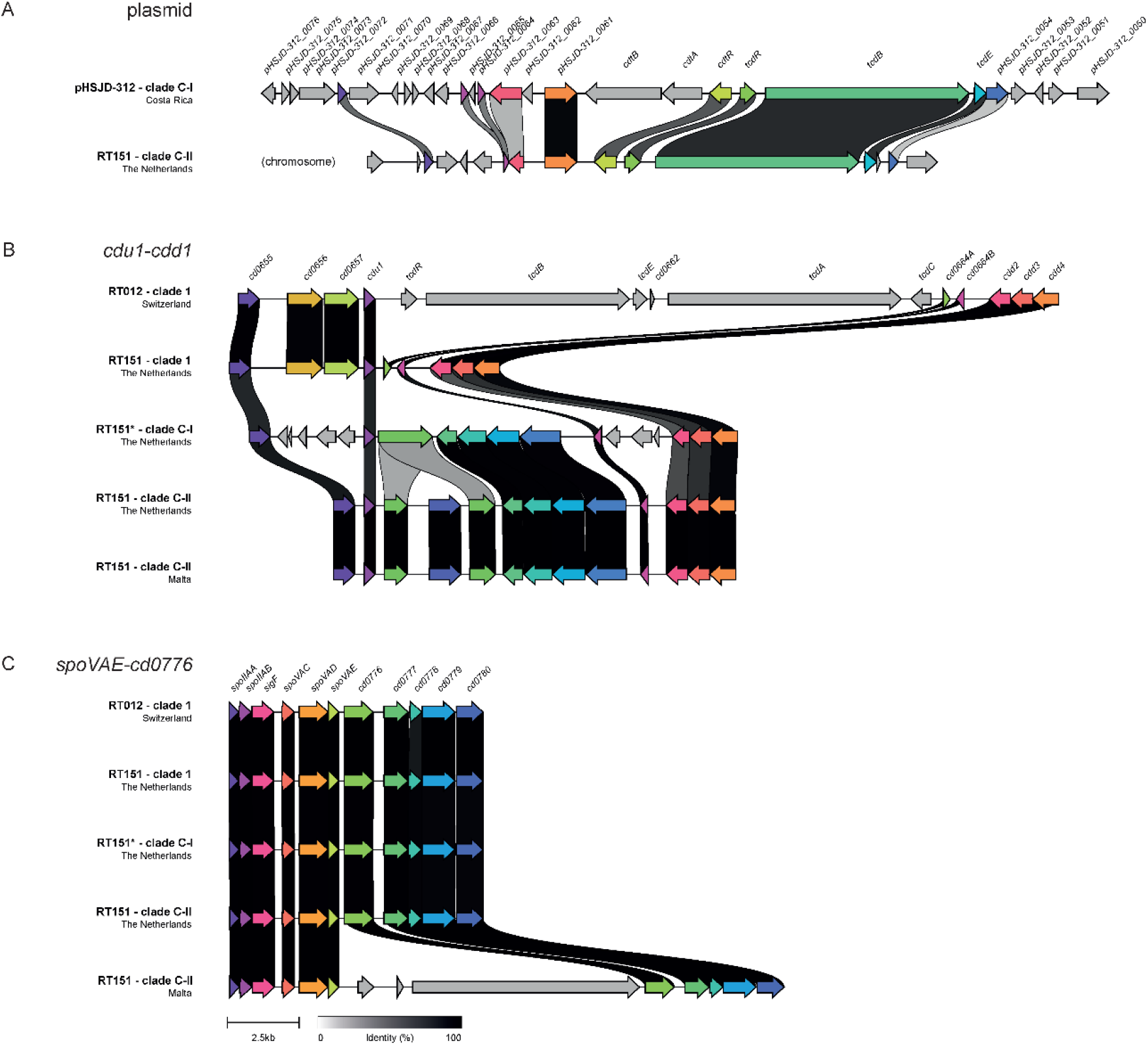
Alignment of previously characterized PaLoc insertion sites for RT151 isolates in comparison with the RT012 reference strain 630(33). **A**. Extrachromosomal element pHSJD-312. **B**. *cdu1-cdd1*. **C**. *spoVAE-cd0776*)(6). RT151 clade 1 is LUMCMM11 0004, RT151 clade C-I is LUMCMM16 0013, and RT151 clade C-II are LUMCMM21 0001/0003 (The Netherlands) and LUMCMM19 2333 (Malta). Figure generated by clinker(43). Colored arrows indicate similar genes; links are drawn between similar genes on neighboring clusters and are shaded based on sequence identity (0% white, 100% black, identity threshold for visualization 0.30).

